# Automated 3D Segmentation of Human Vagus Nerve Fascicles and Epineurium from Micro-Computed Tomography Images Using Anatomy-Aware Neural Networks

**DOI:** 10.1101/2025.06.12.659370

**Authors:** Jichu Zhang, Maryse Lapierre-Landry, Havisha Kalpatthi, Michael W. Jenkins, David L. Wilson, Nicole A. Pelot, Andrew J. Shoffstall

## Abstract

**Objective:** Precise segmentation and quantification of nerve morphology from imaging data are critical for designing effective and selective peripheral nerve stimulation (PNS) therapies. However, prior studies on nerve morphology segmentation suffer from important limitations in both accuracy and efficiency. This study introduces a deep learning approach for robust and automated 3D segmentation of human vagus nerve fascicles and epineurium from high-resolution micro-computed tomography (microCT) images.

**Methods:** We developed a multi-class 3D U-Net to segment fascicles and epineurium that incorporates a novel anatomy-aware loss function to ensure that predictions respect nerve topology. We trained and tested the network using subject-level five-fold cross-validation with 100 microCT sub-volumes (11.4 μm isotropic resolution) from cervical and thoracic vagus nerves stained with phosphotungstic acid from five subjects. We benchmarked the 3D U-Net’s performance against a 2D U-Net using both standard and anatomy-specific segmentation metrics.

**Results:** Our 3D U-Net generated high-quality segmentations (average Dice similarity coefficient: 0.93). Compared to the 2D U-Net, our 3D U-Net yielded significantly better volumetric overlap, boundary delineation, and fascicle-level accuracy. The 3D approach reduced anatomical errors by 2.5-fold, provided more consistent inter-slice boundaries, and improved detection of fascicle splits/merges by nearly 6-fold.

**Significance:** Our automated 3D segmentation pipeline provides anatomically accurate 3D maps of peripheral neural morphology from microCT data. The automation allows for high throughput, and the substantial improvement in segmentation quality and anatomical fidelity enhances the reliability of morphological analysis, vagal pathway mapping, and the implementation of realistic computational models. These advancements provide a foundation for understanding the functional organization of the vagus and other peripheral nerves and optimizing PNS therapies.

## 1 Introduction

Peripheral nerve stimulation (PNS) targeting somatic nerves enables the restoration of movement and sensation following spinal cord injury or limb loss (Taghlabi et al., 2024). PNS targeting autonomic nerves enables treatment of a wide variety of diseases, such as sacral nerve stimulation to restore bladder and bowel function (Assmann et al., 2020) and vagus nerve stimulation (VNS) for numerous indications (Johnson and Wilson, 2018; Pavlov and Tracey, 2022). VNS targets the vagus nerve, which provides extensive sensory and motor innervation between the brainstem and the viscera and critically regulates physiological functions and homeostasis (Neuhuber and Berthoud, 2021). Implanted VNS is FDA-approved to treat epilepsy (Ben-Menachem et al., 1994), depression (Sackeim et al., 2001), obesity (Ikramuddin et al., 2014), and stroke sequelae (Dawson et al., 2021). Ongoing studies are investigating VNS for treating heart failure (Konstam et al., 2019), chronic pain (Shao et al., 2023), and inflammatory disorders (Bonaz et al., 2021). Despite this broad clinical potential, the efficacy of current VNS approaches is limited by their non-selective stimulation of the entire vagus nerve trunk (Fitchett et al., 2021). This lack of selectivity frequently leads to off-target stimulation and adverse effects—including coughing, throat pain, hoarseness, and dyspnea—which compromise clinical outcomes (Ben-Menachem, 2001). Consequently, VNS techniques with improved selectivity are needed to precisely target desired pathways while avoiding those that cause side effects.

Developing selective PNS therapies remains challenging due to an incomplete understanding of the targeted neuroanatomy (Fitchett et al., 2021). Micro-computed tomography (microCT) is an ex vivo nerve imaging technique that can resolve the three-dimensional (3D) structure of fascicles (bundles of fibers) at high resolution (∼5 to 40 μm voxels) with large fields of view (∼10 cm) (Jayaprakash et al., 2023; Thompson et al., 2020; Upadhye et al., 2022). Recent advances in nerve staining for microCT have further improved fascicular contrast and maintained compatibility with histology (Upadhye et al., 2025). Thus, microCT is a key imaging modality for characterizing nerve morphology.

To inform the development of PNS therapies, microCT images must be segmented to extract the nerve (epineurium) and fascicle boundaries. The segmentations enable quantification of the neuroanatomy, including nerve and fascicle diameters (Pelot et al., 2020) and morphological changes due to splits and merges of fascicles (Upadhye et al., 2022). They also enable the analysis of the nerve’s functional organization (Jayaprakash et al., 2023; Thompson et al., 2023). These insights are key to identifying novel points of intervention and designing function- or organ-specific neuromodulation. The segmentations can also serve as inputs for anatomically realistic computational models of PNS: by predicting nerve fiber responses to stimulation, computational models inform the design of electrode geometry, electrode placement, and stimulation parameters to achieve targeted neural responses (Aristovich et al., 2021; Butson et al., 2011; Ciotti et al., 2024; Kent and Grill, 2013; Musselman et al., 2025; Tebcherani et al., 2024; Wongsarnpigoon and Grill, 2010). Therefore, segmentation of nerve morphology is fundamental to quantitative mapping of vagal pathways and drives the design of novel PNS therapies.

Accurate segmentation of nerve morphology is essential, but existing methodologies are limited by throughput and accuracy. Manual or semi-automated segmentation is commonly applied to histological (Pelot et al., 2020; Settell et al., 2020; Verlinden et al., 2016) and microCT (Kronsteiner et al., 2022; Thompson et al., 2023) images but is labor-intensive. Nerve morphology varies substantially between individuals (Brill and Tyler, 2017; Pelot et al., 2020; Upadhye et al., 2022); therefore, large-scale analyses are warranted with accompanying automated segmentation techniques for high throughput, such as convolutional neural networks (CNNs) (Rizwan I Haque and Neubert, 2020). Recent studies have conducted microCT of vagus nerves and used two-dimensional (2D) CNNs for segmentation (Buyukcelik et al., 2023; Jayaprakash et al., 2023). Processing data slice-by-slice, these 2D networks do not leverage the complete spatial context available from microCT data and risk creating inaccurate representations of the vagus nerve’s 3D fascicular connectivity. Further, these prior CNNs treat nerve segmentation as a per-pixel classification task, using generic metrics such as Dice score, which also overlooks the expected anatomical structures (Stewart, 2003). Therefore, the use of existing 2D CNNs to segment microCT images of nerves is unlikely to achieve the segmentation quality demanded by downstream applications— morphological measurements (Brill and Tyler, 2017; Grinberg et al., 2008; Pelot et al., 2020; Schiefer et al., 2012), fascicle tracking (Jayaprakash et al., 2023; Sunderland, 1945; Thompson et al., 2023; Upadhye et al., 2022), and computational modeling (Brill and Tyler, 2011; Musselman et al., 2023; Schiefer et al., 2012)—without extensive manual refinements. Together, these limitations highlight the need for a new automated segmentation approach that is designed and validated based on its ability to create accurate, functionally useful 3D anatomical maps of nerve morphology to inform PNS development.

In this study, we present an enhanced deep learning approach for accurate segmentation of fascicles and epineurium from microCT images of cadaveric human vagus nerves. Our approach features a 3D U-Net CNN that leverages the volumetric context inherent in high-resolution microCT data. We guided network training with a novel anatomy-aware loss function that incorporates structural constraints of the nerve. We benchmarked the segmentation accuracy of our 3D U-Net against a published 2D U-Net (Buyukcelik et al., 2023) using five-fold cross-validation with both standard and anatomy-aware metrics. Our 3D approach achieved significantly better spatial overlap, boundary delineation, and detection of individual fascicles. The resulting 3D segmentations also showed improved preservation of fascicle connectivity, fewer anatomical errors, and more consistent boundaries across slices. By overcoming key limitations of prior techniques, our segmentation pipeline provides a high-throughput tool for generating realistic 3D anatomical maps of the vagus nerve and is adaptable to other PNS targets. These improved segmentations facilitate analyses that can advance our understanding of vagal neuroanatomy and accelerate the development of next-generation VNS therapies.

## 2 Methods

All analyses were performed in R (version 4.3.3) with rstatix (Kassambara, 2023) for statistical tests and ggplot2 (Wickham, 2016) for plotting.

### 2.1 Tissue Collection and Preparation

We collected human vagus nerves from five embalmed cadavers (2 female, 3 male; 57–81 years) donated to the Anatomical Gift Program at Case Western Reserve University (Table S1). This study received non-human subject exemption from the Case Western Reserve University Institutional Review Board.

We dissected the vagus nerve bilaterally from the inferior border of the jugular foramen to the superior end of the esophageal plexus. We mounted the excised nerves on custom acrylic boards (3 cm wide, up to 9 cm long); the nerve tissue on each board was termed a “sample”. We stained nerve samples with 3% phosphotungstic acid (PTA; HT152-250ML, Sigma-Aldrich, St. Louis, MO) for 24 hours and gently agitated at 50 rpm using an orbital shaker (SI-M1500, Scientific Industries, Bohemia, NY) (Upadhye et al., 2025). After staining, we covered the samples with gauze saturated with 1X phosphate-buffered saline (PBS; BP399-1, Fisher Scientific, Hampton, NH) and stored them in sealed containers at 4°C for 0.5 to 4 days.

### 2.2 MicroCT Image Acquisition

We loaded the nerve samples mounted on their acrylic boards into a cylindrical sample tube (34 mm diameter × 110 mm height, U50825, Scanco Medical AG, Brüttisellen, Switzerland). We performed microCT scanning using a μCT 100 cabinet scanner (Scanco Medical AG) with the following parameters: X-ray voltage 55 kV, current 145 mA, integration time 500 ms, and 0.5 mm aluminum filter. These parameters were selected to balance image quality with scan duration, reducing the risk of tissue dehydration (Upadhye et al., 2025). We scanned all samples with a cross-sectional field of view of 35.2 mm diameter at 11.4 μm isotropic resolution. Scanning each tube (containing two nerve samples) required ∼14 hours. We reconstructed scans using the manufacturer’s proprietary software and exported as series of 16-bit DICOM images (3072 × 3072 pixels). For more efficient file storage and analysis, we compiled the DICOM image data for each sample into 3D OME-Zarr arrays (Moore et al., 2023) at the original resolution.

### 2.3 Ground Truth Annotation

We created a dataset with 100 non-overlapping microCT sub-volumes evenly distributed across the five cadavers. Each sub-volume was 64 × 1536 × 3072 voxels (*z, y, x*) in size with an isotropic resolution of 11.4 μm; the *x* and *y* dimensions captured the full nerve cross section in all slices, and the *z* dimension spanned 0.73 mm per sub-volume. We selected these sub-volumes from representative cervical and thoracic regions containing fascicle splits and merges, and we avoided areas with excessive tissue damage. We exported each sub-volume at the original resolution as a 16-bit TIFF stack for ground truth annotation.

We manually segmented fascicle and epineurium boundaries from 100 microCT images from five cadaveric subjects to generate ground truth annotations for training the 3D U-Net (Figure 1a). We manually traced fascicle boundaries every 10 cross sections in a sub-volume using 3D Slicer (version 5.6.1) (Fedorov et al., 2012). We interpolated between manually segmented slices using the “Fill between slices” function. We reviewed and edited the segmentations at each fascicle split or merge.

**Figure 1.**
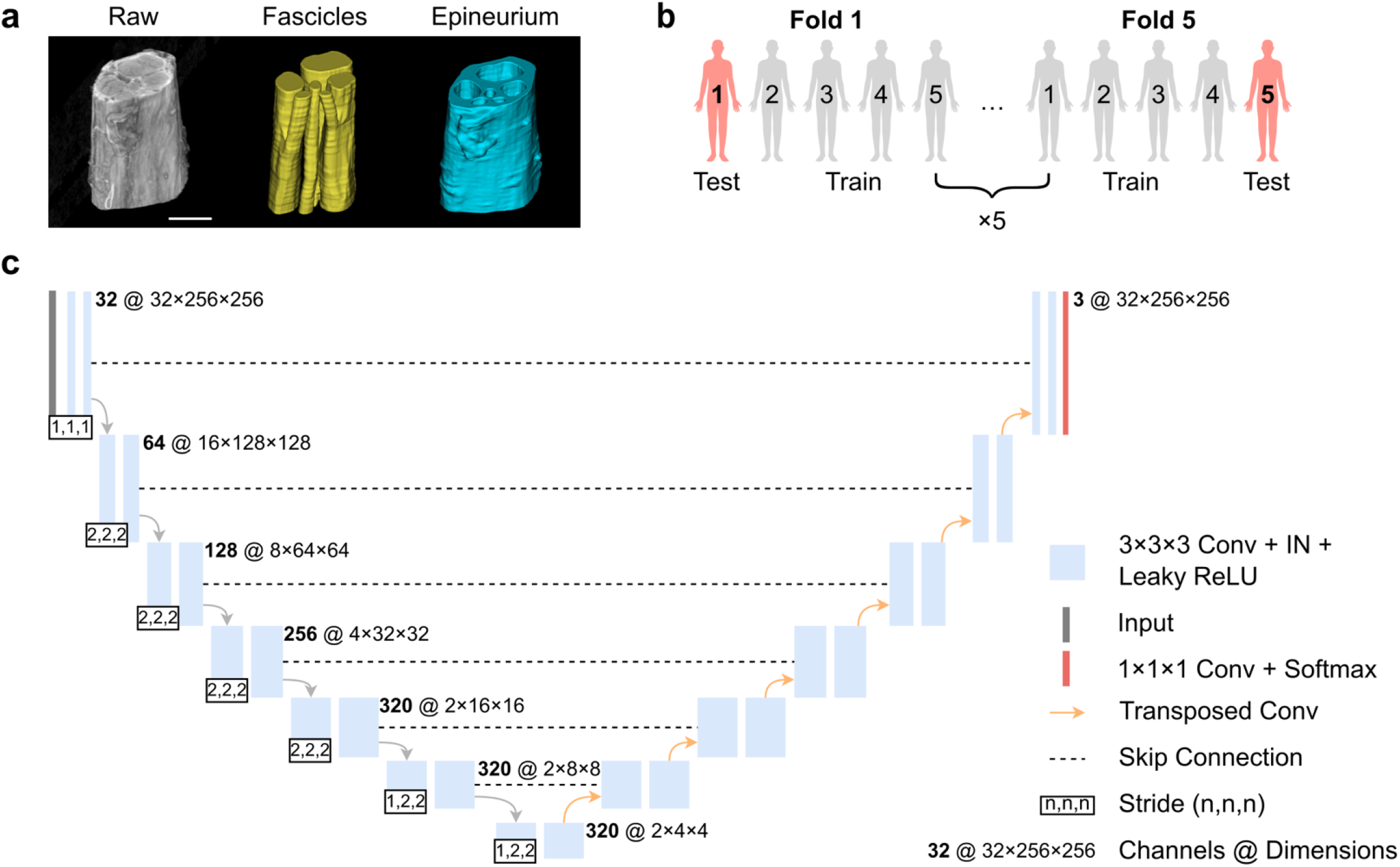
Ground truth annotation, cross-validation strategy, and 3D U-Net architecture. **(a)** Example of a raw microCT sub-volume and corresponding manual segmentation of fascicles and epineurium. Scale bar is 500 μm. **(b)** The five-fold leave-one-out cross-validation approach used for training and evaluation. Each iteration used four subjects (80 images) for training and one subject (20 images) for testing. **(c)** Multi-class 3D U-Net architecture for fascicle and epineurium segmentation. Conv, convolution; IN, instance normalization; ReLU, rectified linear unit.

We segmented the epineurium semi-automatically using the Segment Anything for Microscopy package (Archit et al., 2025). We first applied the built-in vision transformer Base (ViT-B) model to interactively segment the epineurium every 16 cross sections in a sub-volume followed by manual corrections. Then, we used the “Automatic tracking” function to propagate these “seed” masks across the entire image stack with the following parameters: intersection-of-union (IoU) threshold = 0.5, motion smoothing factor = 0.5, bounding box extension = 0.05.

We combined the fascicle and epineurium segmentations into an 8-bit TIFF stack at the original image resolution. Each voxel was classified as fascicle, epineurium, or background; fascicle labels overwrote epineurium in cases of overlap. We validated each segmentation against the raw microCT sub-volume, verifying boundary accuracy, fascicle splits and merges, and structural continuity across slices.

### 2.4 Network and Loss Function

#### 2.4.1 Network Architecture

Our neural network is based on published 3D U-Net architectures (Çiçek et al., 2016; Ronneberger et al., 2015) and was implemented using the nnU-Net framework (Isensee et al., 2021), which automatically configured the architecture and hyperparameters based on properties of the input dataset. The network processed single-channel grayscale microCT volumes and output three-class segmentation maps wherein each voxel was identified as background, fascicle, or epineurium.

The network featured a 7-stage encoder-decoder with progressive feature dimensions of 32, 64, 128, 256, 320, 320, and 320 (Figure 1c). Each stage had two 3 × 3 × 3 convolutions with varying downsampling strides: (1,1,1) for stage 1, (2,2,2) for stages 2 to 5, and (1,2,2) for stages 6 and 7. The decoder path mirrored the encoder structure with skip connections from corresponding stages, maintaining consistent kernel sizes and strides except at the bottleneck layer. We used instance normalization and leaky ReLU activation throughout the network without dropout.

We adopted the published implementation for 3D nnU-Net (Isensee et al., 2021) with Python 3.10 and PyTorch 2.1.2. All training and evaluation were performed on an NVIDIA RTX A6000 GPU (48 GB memory; Santa Clara, CA) with CUDA 12.1 using 16-bit automatic mixed precision.

#### 2.4.2 Anatomy-Aware Loss Function

To ensure anatomically plausible segmentation of nerve structures, we adopted an anatomy-aware module based on the topological interaction framework (Gupta et al., 2022). The module enforced two anatomical constraints: (1) fascicles must be completely enclosed by epineurium, and (2) fascicles cannot directly contact background voxels. Using 3D convolution operations based on 26-neighbor voxel connectivity, the module identified voxels where these anatomical rules were violated (Figure 2a). The resulting critical voxel map *V* highlighted regions that required additional supervision during training.

**Figure 2.**
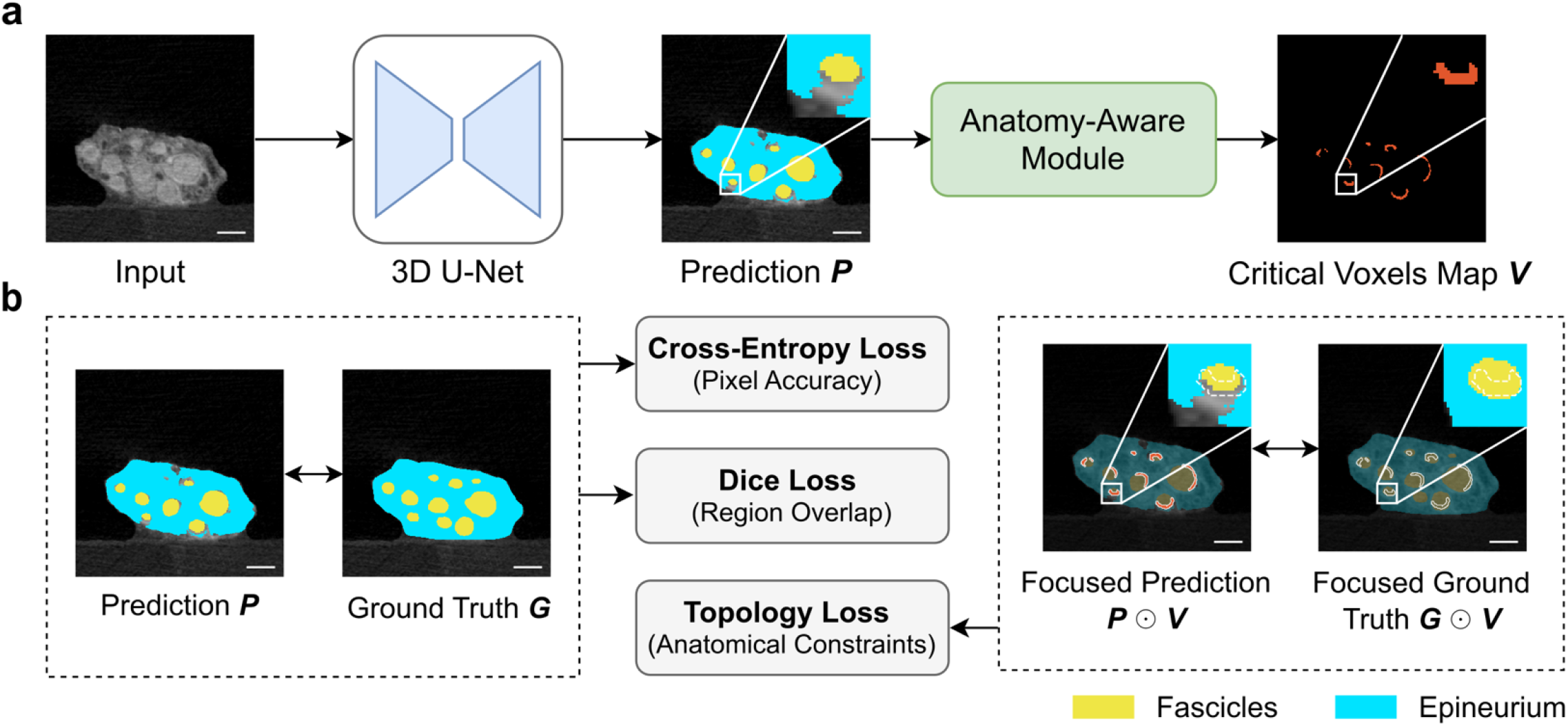
Overview of the anatomy-aware network training and compound loss computation. **(a)** Identification of voxels in the automated segmentation that do not meet anatomical constraints, i.e., fascicles must be completely enclosed by epineurium and fascicles cannot directly contact background. The anatomy-aware module identifies regions in the 3D U-Net prediction where anatomical constraints are violated and generates a mask of critical voxels *V* that highlights areas requiring focused supervision during training. **(b)** Computation of the compound loss function. The network is trained using three loss terms: cross-entropy loss (*L*_CE_) for pixel-wise accuracy, Dice loss (*L*_Dice_) for region overlap, and topology loss (*L*_Topo_) for a priori anatomical knowledge. The topology loss is computed by applying cross-entropy loss only to regions identified in the critical voxels map *V*. ⊙ denotes the Hadamard product. Scale bars are 500 μm.

We incorporated anatomical constraints into the network training through a compound loss function (Figure 2b) with three components: cross-entropy loss (*L*_CE_) for pixel-wise classification accuracy, Dice loss (*L*_Dice_) for optimizing region overlap, and topology loss (*L*_Topo_) for enforcing anatomical constraints. The topology loss was computed by applying cross-entropy loss specifically to regions identified in the critical voxels map *V*:

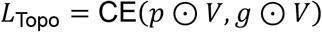

where *p* is the predicted segmentation, *g* is the ground truth, and ⊙ is the Hadamard product. The final loss function combines all components:

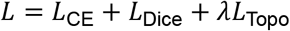

with λ = 1 × 10^6^ to weight the topological component of the 3D prediction (Gupta et al., 2022). This compound loss function ensured both global segmentation quality and appropriate anatomical relationships between neural structures.

### 2.5 Network Training and Inference

We implemented a five-fold leave-one-out cross-validation strategy (Figure 1b). The dataset of 100 annotated microCT volumes was divided at the subject level to prevent data leakage. In each iteration, we used 80 volumes from four subjects for training and 20 from the remaining subject were used for testing. This subject-wise partitioning allowed us to evaluate the network’s generalized performance across different anatomical variations and quality of tissue, staining, and imaging.

Our 3D U-Net was trained using patches of size 32 × 256 × 256 voxels randomly sampled from the full-resolution volumes (11.4 × 11.4 × 11.4 μm^3^). We ensured that 33% of the training patches contained at least one foreground class (i.e., fascicle or epineurium). Before training, all input images were normalized by clipping the original 16-bit intensities to [0,32767] and rescaling to [0,1]. To prevent overfitting, we applied an extensive set of data augmentation techniques during training, including random rotation, random noise, and contrast adjustments; see Table S2 in Supplementary Note 2 for the complete list of data augmentations with parameter ranges and probabilities.

We trained the 3D U-Net for 500 epochs with a batch size of 4 patches, where each epoch consisted of 250 mini-batch iterations (see Figure S1 in Supplementary Note 2 for example training curves). We used stochastic gradient descent with Nesterov momentum (*μ* = 0.99) as the optimizer. The initial learning rate was set to 0.01 and followed a polynomial decay schedule, where the learning rate at each epoch was reduced by multiplying it by (1 − *e*/*e*_max_)^0.9^. Here, *e* represented the current epoch number and *e*_max_ was the total number of epochs. After each training epoch, we performed in-training validation on 50 patches randomly extracted from the validation set rather than full-size images, allowing for efficient training without the computational overhead of full image inference.

During full image inference, we extracted overlapping patches at a regular interval of 8 × 64 × 64 voxels (25% overlap in all dimensions). We merged the overlapping patches using a sliding window approach with Gaussian blending (σ = 0.125) weighted by patch centers.

### 2.6 3D U-Net vs. 2D U-Net Comparison

We compared the performance of our 3D U-Net to a published 2D U-Net that was also implemented to segment the fascicular morphology of human vagus nerves from microCT images (Buyukcelik et al., 2023) (Figure S2). We optimized the 2D U-Net’s hyperparameters and trained it on the same dataset using identical cross-validation splits, preprocessing steps, and data augmentation techniques. For the 2D U-Net, we used the Adam optimizer with a constant learning rate of 0.0001 and a batch size of 8 patches of size 512 × 512 pixels. The sampling strategy ensured that 67% of the patches contained at least one foreground class (i.e., fascicle or epineurium). We performed a grid hyperparameter search to determine the optimal learning rate, batch size, and patch size. For in-training validation, an 80/20 split was performed at the volume level (instead of slice-wise) to prevent data leakage from similar, adjacent cross sections.

During inference, the 2D U-Net processed raw microCT volumes slice-by-slice with 512 × 512 patches extracted at a regular interval of 128 × 128 pixels (25% overlap in both dimensions). We merged the patches using the same sliding window approach as the 3D U-Net. We stacked the slice-wise predictions into 3D volumes for evaluation and visualization.

We compared the networks using paired, two-sided Wilcoxon signed-rank tests (α = 0.05) for each metric. We reported 95% confidence intervals derived from bootstrap resampling (n = 1000) and *p*-values with a Bonferroni correction for multiple comparisons.

### 2.7 Ablation Study

To evaluate the contribution of the anatomy-aware component of the loss function, we performed an ablation study by training an identical 3D U-Net architecture with only the conventional loss terms—Dice loss (*L*_Dice_) and cross-entropy loss (*L*_CE_). The same training protocol, data splits, augmentation techniques, and optimization hyperparameters were maintained for both 3D U-Nets.

### 2.8 Evaluation Metrics

We evaluated the performance of the U-Nets using a set of quantitative metrics. *P* denotes the predicted segmentation (output from the network), and *G* denotes the ground truth segmentation (manual annotations). For each test image, all metrics were calculated separately for fascicle and epineurium classes (where applicable).

#### 2.8.1 Segmentation Accuracy

##### Dice Similarity Coefficient

The Dice similarity coefficient (DSC) (Dice, 1945) measures the volumetric overlap between the predicted and ground truth segmentations, with values ranging from 0 (no overlap) to 1 (perfect overlap):

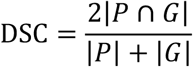

where |*P* ∩ *G*| represents the number of true positive voxels, and |*P*| and |*G*| represent the total number of voxels in the predicted and ground truth segmentations, respectively.

##### Surface Dice Similarity Coefficient

The surface DSC (Nikolov et al., 2021) quantifies boundary accuracy by measuring the overlap between predicted and ground truth surfaces within a specified tolerance, with values ranging from 0 (no surface overlap) to 1 (perfect surface overlap). For a solid label *L* with surface *S*(*L*), its border region *B* is defined as:

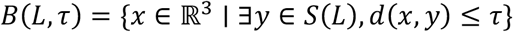

where *d*(·) denotes the Euclidean distance and τ is a distance tolerance, which we set to one voxel. The surface DSC is then calculated as:

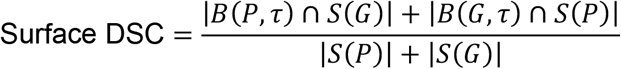

where *S*(*P*) is the surface of the predicted segmentation, *S*(*G*) is the surface of the ground truth segmentation, and the intersection terms represent the portions of each surface that fall within the border region of the other.

##### Average Symmetric Surface Distance (ASSD)

The ASSD (Heimann et al., 2009) measures the average physical distance between the surfaces of the predicted and ground truth segmentations, indicating the magnitude of segmentation errors. A lower ASSD value indicates better agreement between the predicted and ground truth surfaces:

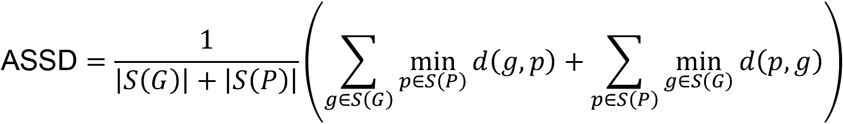

where *d*(·) denotes the Euclidean distance, *g* is a point in ground truth segmentation, and *p* is a point in predicted segmentation.

#### 2.8.2 Fascicle Detection Accuracy

In addition to standard segmentation metrics, we evaluated the network’s ability to correctly identify fascicle instances in cross sections. We defined binary masks of fascicles (*F*) where voxels belonging to fascicle tissue were assigned a value of 1 and all other voxels were assigned a value of 0. We identified individual fascicle instances by applying an 8-connectivity connected components algorithm to the fascicle masks. We matched ground truth (*F*_*G,i*_) and predicted (*F*_*P,j*_) instances using intersection over union (IoU):

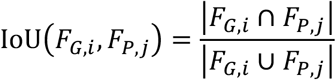

For a given IoU threshold *t*, we identified the matching pairs (i.e., true positives) of fascicle instances where IoU(*F*_*G,i*_, *F*_*P,j*_) ≥ *t*. Based on the matches, we calculated the fascicle *F*_1_ score at threshold *t* as:

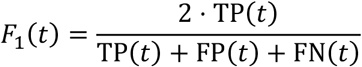

where TP(*t*), FP(*t*), and FN(*t*) are the number of true positives, false positives, and false negatives at threshold *t*, respectively.

We evaluated the *F*_1_ score across IoU thresholds ranging from 0.50 to 0.95, with increments of 0.05. The *F*_1_ score balances precision and recall, where a prediction is considered a true positive if its IoU with a ground truth fascicle exceeds the threshold. Using a threshold of *t* ≥ 0.5 ensured that each ground truth fascicle had at most one match in the predictions, and vice versa.

At a threshold of *t* = 0.7, we quantified the fraction of fascicles that were over-segmented (i.e., one ground truth fascicle was split into multiple predicted fascicles) or under-segmented (i.e., multiple ground truth fascicles were merged into a single predicted fascicle). We also measured the fraction of missed fascicles (i.e., false negatives) by cross-sectional area (Buyukcelik et al., 2023): tiny (< 0.02 mm^2^), small (0.02–0.09 mm^2^), medium (0.09– 0.3 mm^2^), and large (> 0.3 mm^2^). The four categories corresponded to effective circular diameters of < 0.16 mm, 0.16–0.34 mm, 0.34–0.62 mm, and > 0.62 mm, respectively. We defined the effective circular diameter as the diameter of a circle with the same area as the original fascicle segmentation.

Over-segmentation, under-segmentation, and missed fascicle rates were calculated as the proportion of affected fascicles relative to the total number of ground truth fascicles. These metrics were computed at 8-slice intervals (∼0.1 mm) and averaged across a test volume.

#### 2.8.3 Anatomical Accuracy

In addition to standard segmentation metrics and object-level precision, we also evaluated the anatomical coherence of the network’s predictions based on the preservation of 3D fascicle connectivity and the rate of topological violations.

##### Centerline Dice

The centerline Dice score (clDice) (Shit et al., 2021) measures the network’s ability to preserve the correct anatomical connectivity of fascicles. To calculate this similarity metric, we computed the topology precision (*T*_prec_) and sensitivity (*T*_sens_) as:

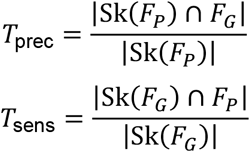

where Sk(*F*_*P*_) and Sk(*F*_*G*_) are the skeletonized predicted and ground truth fascicle masks, respectively, generated using the skeletonize function in the Python scikit-image package (Walt et al., 2014). The clDice is then calculated as:

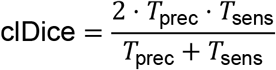

with values ranging from 0 (no topology preservation) to 1 (perfect topology preservation).

##### Anatomical Error Rate

We quantified the anatomical quality of neural tissue predictions by measuring the frequency of structural abnormalities, as detailed in Supplement 8.3. Two types of errors were identified: (1) exposed fascicle voxels that directly contacted background voxels, and (2) abrupt transitions between neural structures along the nerve. The anatomical error rate was calculated as the ratio of anomalous voxels to total foreground voxels (i.e., fascicle and epineurium).

##### Inter-Slice Boundary Consistency

The inter-slice boundary consistency quantifies the stability of predicted contours along the nerve, i.e., the lack of inter-slice jitter of a given boundary. For each pair of adjacent slices *z* and *z* + 1, we computed the boundary *F*_1_ score (BF score) (Csurka et al., 2013) with single-pixel tolerance. Let *B*_*z*_ and *B*_*z*+1_ be the boundary regions (one-pixel-wide contours) of the predicted mask in slices *z* and *z* + 1, respectively. We calculated the boundary precision (*P*_*z,z*+1_) and recall (*R*_*z,z*+1_) as:

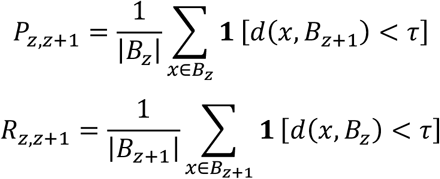

where *d*(·) denotes the Euclidean distance, **1**[·] is the indicator function (1 if the condition is true, 0 otherwise), and τ = 1 is the distance tolerance in pixels. The BF score between slices *z* and *z* + 1 is then calculated as:

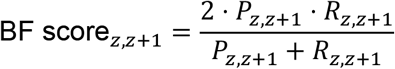

with values ranging from 0 (no boundary consistency) to 1 (perfect boundary consistency).

##### Fascicle Split/Merge Detection

Fascicles split and merge along the nerve (Upadhye et al., 2022). We evaluated the model’s ability to detect these structural changes by analyzing the rates of split/merge events in predictions versus ground truth.

We identified fascicle split/merge locations in ground truth and network prediction by analyzing the division and fusing of fascicle boundaries (Upadhye et al., 2022). For each image, we calculated the rate of split or merge events per millimeter as *R*_split_ = *N*_split_/*L* and *R*_merge_ = *N*_merge_/*L*, where *N*_split_ and *N*_merge_ are the number of detected split and merge events, respectively, and *L* is the length of the nerve segment (along the z axis) in millimeters. The deviation from the ground truth event rate was quantified as *ΔR* = *R*_*P*_ − *R*_*G*_, where *R*_*P*_ and *R*_*G*_ are the predicted and ground truth event rates, respectively. For both split and merge events, we computed:

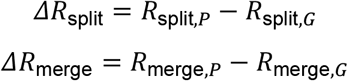

The split/merge event rate deviation was quantified as the average absolute difference between predicted and ground truth rates:

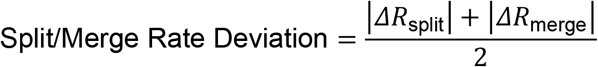

with lower values indicating better split/merge detection accuracy.

## 3 Results

We developed an automated, deep learning-based approach to segment volumetric microCT images of cadaveric human vagus nerves by delineating the nerve boundary and 3D fascicular structure. We trained a 3D U-Net and evaluated its performance through segmentation metrics and by assessing its consistency with expected anatomical structures.

### 3.1 Segmentation Accuracy

We implemented a 3D U-Net to segment vagus nerve fascicles and epineurium in microCT images and evaluated its performance using five-fold cross-validation. The 3D U-Net outperformed the 2D U-Net in both spatial overlap and boundary accuracy metrics (Figure 3). Segmentation maps from three representative nerve samples, selected from the test set of their respective folds, showed that the 3D U-Net produced clearer, smoother fascicle boundaries with fewer artifacts compared to the 2D U-Net segmentations (Figure 3a).

**Figure 3.**
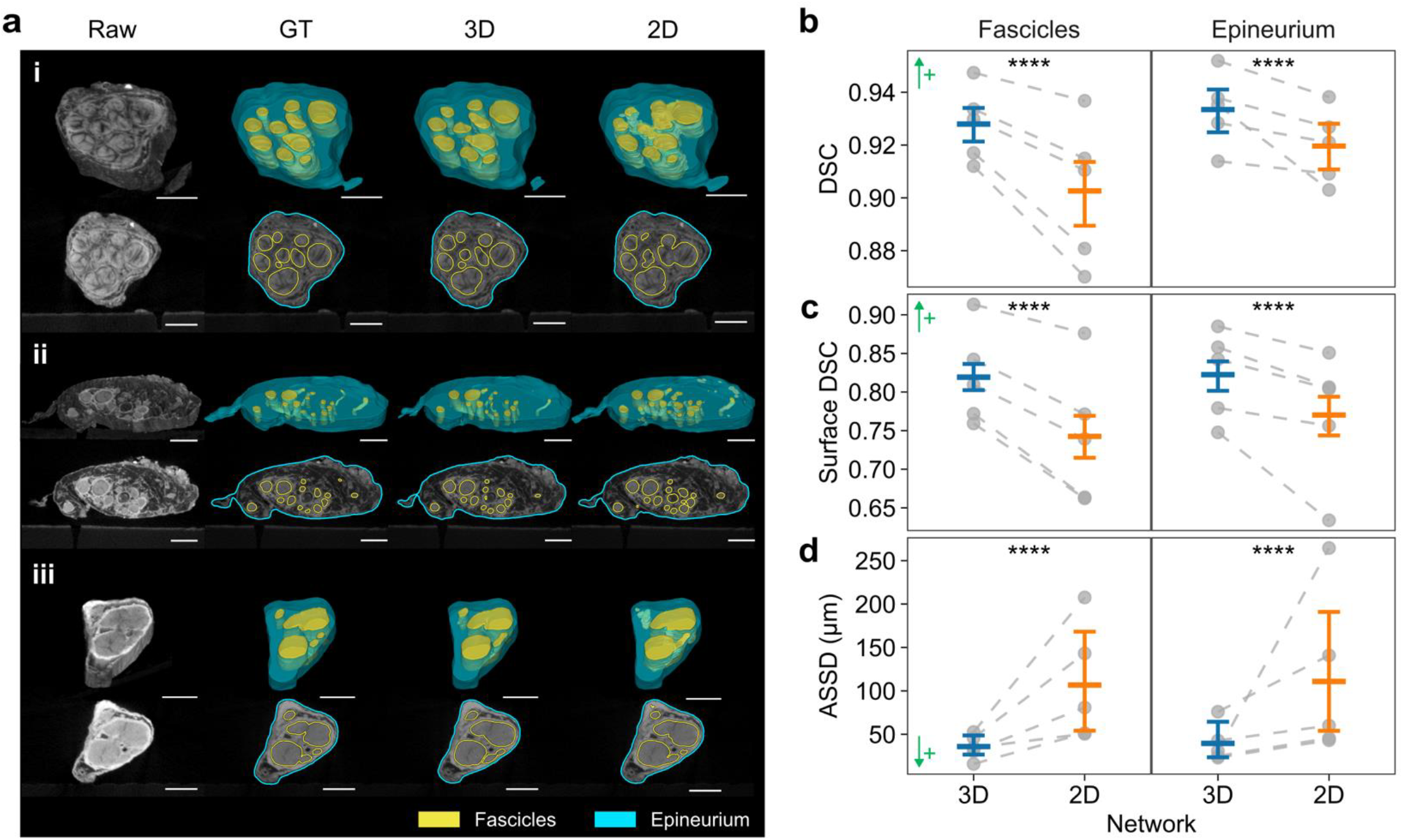
3D U-Net yields more accurate automated segmentations of vagus nerve fascicles and epineurium from microCT compared to a 2D U-Net. **(a)** Representative results comparing 3D and 2D U-Nets for three nerve samples ranked by 3D U-Net Dice similarity coefficient (DSC) at (i) high (90th percentile), (ii) medium (50th percentile), and (iii) low (20th percentile) performance. Images show 3D rendered volumes (top row of each subpanel) and corresponding middle cross sections (bottom row of each subpanel) of raw microCT, ground truth (GT), and predictions from 3D and 2D U-Nets. Scale bars are 1 mm. **(b–d)** Segmentation metrics for each class (fascicles or epineurium) across five cross-validation folds (*N* = 100 images total). Plots show mean values with 95% confidence intervals; individual fold means are shown as gray dots connected by dashed lines. Performance metrics include: **(b)** DSC, **(c)** surface DSC (single-pixel tolerance), and **(d)** average symmetric surface distance (ASSD, in μm). ↑+, higher is better; ↓+, lower is better. ****, *p* < 0.0001.

Quantitatively, the 3D U-Net demonstrated superior segmentation performance compared to the 2D U-Net across multiple metrics. For volumetric overlap, it achieved higher DSCs for both fascicles (0.928 vs. 0.903, *p* < 0.0001) and epineurium (0.933 vs. 0.920, *p* < 0.0001; Figure 3b). Similarly, the 3D U-Net improved boundary delineation, as shown by higher surface DSC values under single-pixel tolerance for fascicles (0.819 vs. 0.743, *p* < 0.0001) and epineurium (0.822 vs. 0.770, *p* < 0.0001; Figure 3c). The 3D U-Net significantly reduced absolute boundary error, as indicated by smaller ASSD values for fascicles (35.9 μm vs. 107 μm, *p* < 0.0001) and epineurium (39.5 μm vs. 111 μm, *p* < 0.0001; Figure 3d). Additional voxel-wise metrics further confirmed the improved performance of 3D U-Net across IoU, sensitivity, and specificity measurements (Table S3 in Supplementary Note 4).

### 3.2 Detection of Fascicle Instances in Cross Sections

We compared the accuracy of the 3D and 2D U-Nets in identifying individual fascicle in cross sections by matching predicted instances with the ground truth using IoU overlap and by calculating the fascicle *F*_1_ score at different levels of stringency to identify a “match”.

A qualitative comparison of two example nerve samples revealed clear differences, where the 3D U-Net identified more fascicle instances and produced fewer segmentation errors than the 2D U-Net (Figure 4a). The quantitative performance was also better for the 3D U-Net, which had higher fascicle *F*_1_ scores across IoU thresholds from 0.5 to 0.9 (Figure 4b). At the IoU threshold = 0.7, the 3D U-Net produced fewer under-segmentation errors (i.e., merging adjacent fascicles into one) than the 2D U-Net (3.94% vs. 4.62%, *p* < 0.01), while both showed similar over-segmentation rates (*p* = 0.66) (Figure 4c).

**Figure 4.**
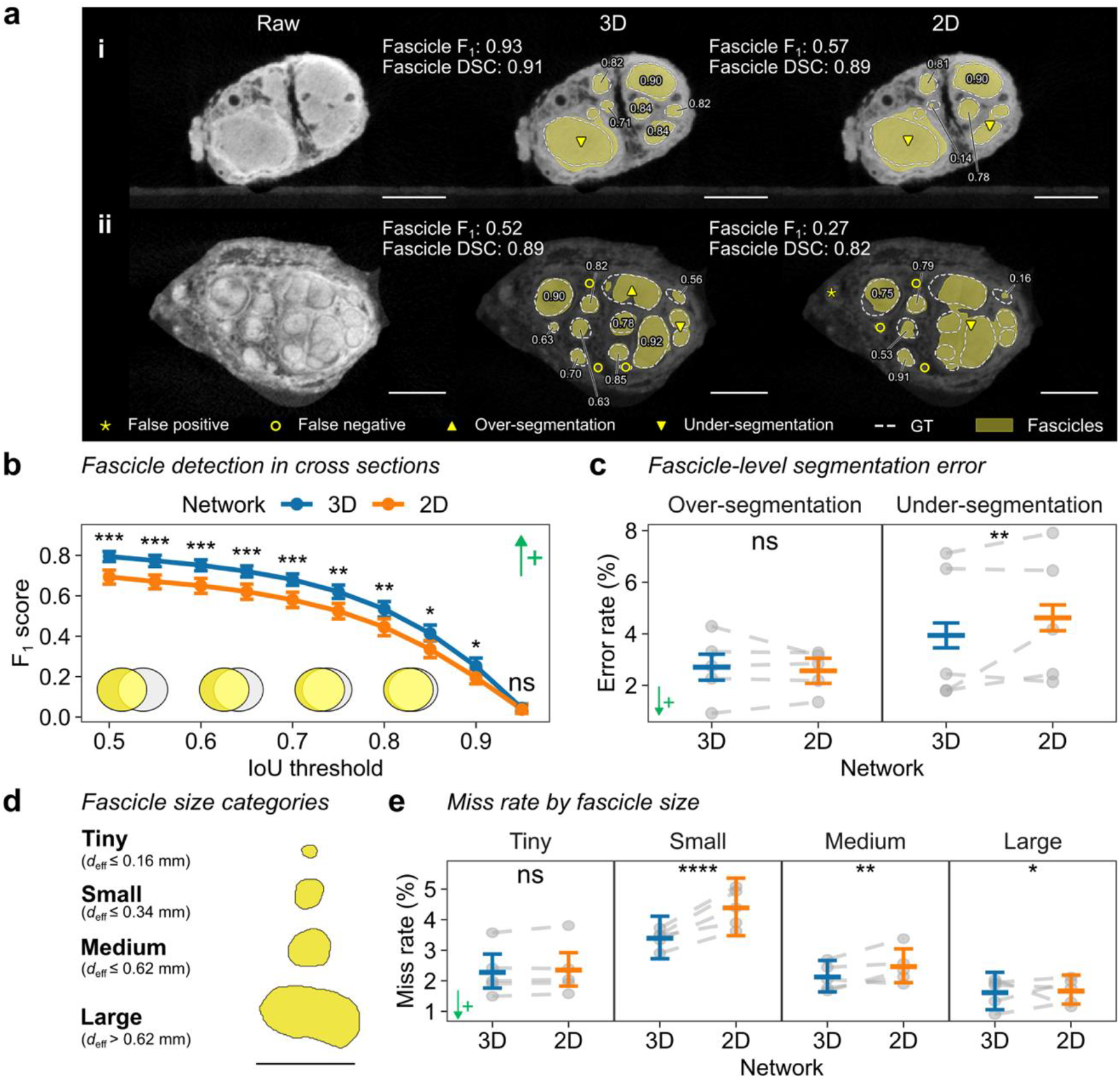
3D U-Net improves detection of individual fascicles in nerve cross sections. **(a)** MicroCT cross sections and fascicle segmentations by 3D and 2D U-Nets for two nerve samples ranked by 3D U-Net’s fascicle **F**_**1**_ score at (i) high (90th percentile) and (ii) low (20th percentile) performance. Fascicle **F**_**1**_ score (at intersection-over-union [IoU] threshold = 0.7) and Dice similarity coefficient (DSC) are labeled. Scale bars are 1 mm. Segmented fascicles (yellow) are overlaid with ground truth fascicles (GT, white dashed lines) with per-fascicle IoU values (omitted for false positives/negatives). **(b)** Mean fascicle **F**_**1**_ score with 95% confidence interval (CI) at various IoU thresholds for 3D and 2D U-Nets. Higher IoU means stricter overlap criteria for matching. **(c)** Comparison of per-fascicle over-segmentation (left) and under-segmentation (right) rates. **(d)** Fascicle size classification into four categories (Buyukcelik et al., 2023) (see distribution in Figure S3 in Supplementary Note 5). *d*_eff_, effective circular diameter. Scale bar is 1 mm. **(e)** Percentage of missed fascicles. In **(c, e)**, plots show mean values with 95% CIs; individual fold means are shown as gray dots connected by dashed lines. ↑+, higher is better; ↓+, lower is better. ****, *p* < 0.0001; ***, *p* < 0.001; **, *p* < 0.01; *, *p* < 0.05; ns, not significant.

A size-based fascicle detection analysis indicated that the 3D U-Net more effectively captured small (miss rate: 3.39% vs. 4.39%, *p* < 0.001) and medium (miss rate: 2.16% vs. 2.42%, *p* < 0.01) fascicles compared to the 2D U-Net (Figure 4d–e). These two size categories constituted 66% of all fascicles in our dataset (Figure S3 in Supplementary Note 5). Both approaches performed similarly for tiny (*p* = 0.059) and large (*p* = 0.047) fascicles. For successfully matched fascicles, the 3D U-Net achieved better instance-level segmentation accuracy, with higher IoU and lower Hausdorff distance values for small and medium fascicles (Table S4 in Supplementary Note 5).

### 3.3 Anatomical Accuracy

Visual comparisons showed that the 3D U-Net maintained continuous fascicular paths, while the 2D U-Net produced both false connections and disconnected paths (Figure 5a). Quantitatively, the 3D U-Net yielded higher clDice scores than the 2D U-Net (0.90 vs. 0.84, *p* < 0.0001; Figure 5b), confirming greater accuracy in the predicted fascicle connectivity. Further, the 3D approach lowered anatomical error rates by ∼2.5-fold compared to the 2D U-Net (0.65% vs. 1.60%, *p* < 0.0001; Figure 5c, d), representing on average ∼9000 fewer erroneous voxels per test image. Ablation experiments—wherein we trained the 3D U-Net without the anatomy-aware loss—confirmed that a priori anatomical knowledge enhanced our 3D U-Net’s ability to preserve nerve structure compared to a baseline 3D architecture (Table S5 in Supplementary Note 6).

**Figure 5.**
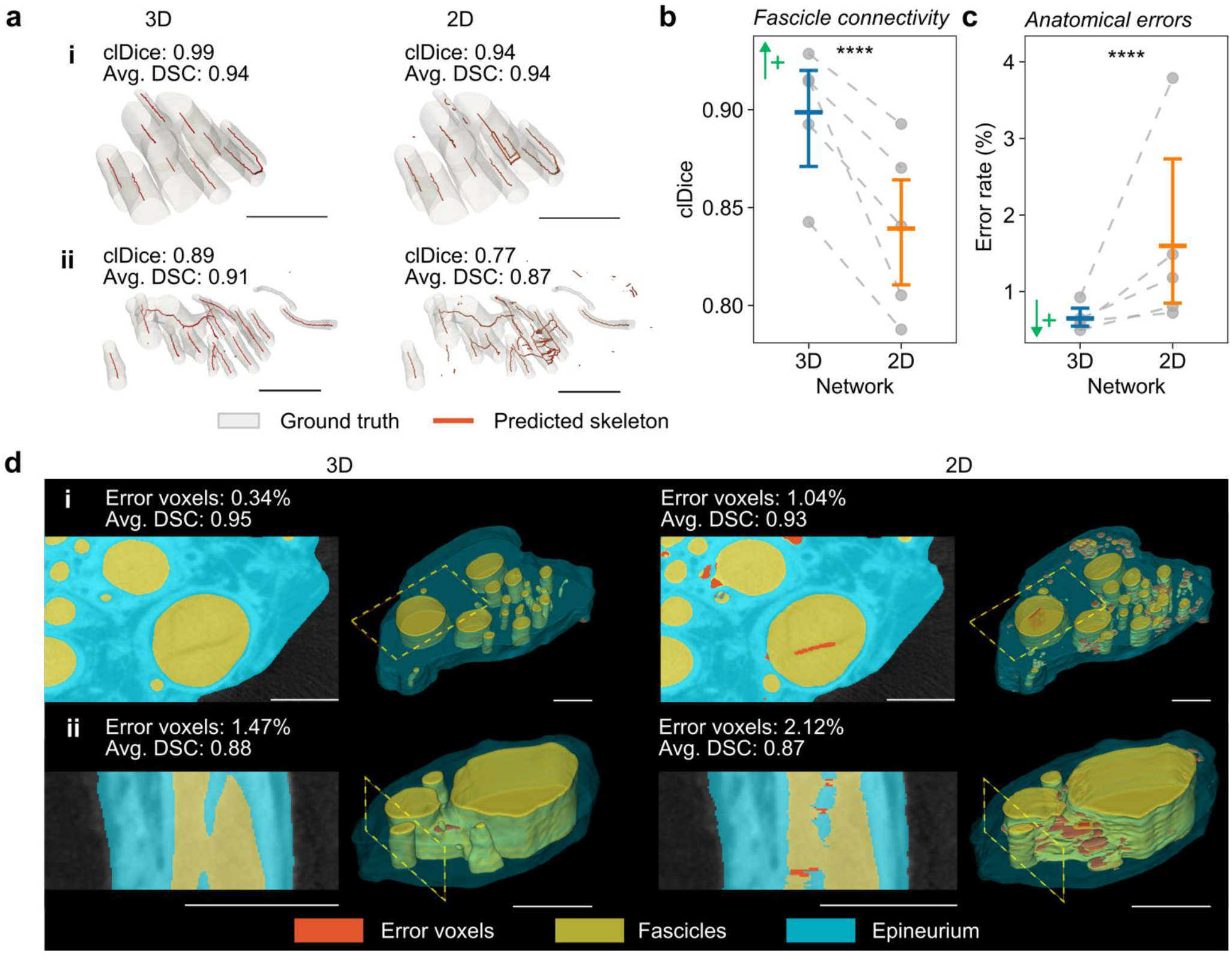
3D U-Net preserves fascicle connectivity and reduces anatomical violations. **(a)** Predicted fascicle skeletons (red) vs. ground truth (GT, gray) for two examples ranked by 3D U-Net centerline Dice (clDice) score at (i) high (90th percentile) and (ii) medium (50th percentile) performance. clDice and average Dice similarity coefficient (DSC) values are labeled. Quantitative comparisons of **(b)** fascicle connectivity measured by clDice scores and **(c)** anatomical error rates (%) between 3D and 2D U-Nets. Plots show mean values with 95% confidence intervals; individual fold means are shown as gray dots connected by dashed lines. **(d)** Visualization of errors violating expected nerve anatomy (red voxels) for two samples ranked by 3D U-Net error rate at (i) low (20th percentile) and (ii) high (90th percentile) violation levels. Images show cross-sectional and 3D views, illustrating typical errors like broken and discontinuous fascicles. Cross section locations are marked by yellow dashed boxes on the 3D volumes. Voxel error rates and average DSC of example images are labeled. Scale bars are 1 mm. ↑+, higher is better; ↓+, lower is better. ****, *p* < 0.0001.

The 3D U-Net showed greater inter-slice consistency than the 2D U-Net or the GT annotation (Figure 6a, c, d). The 3D U-Net generated fascicle segmentations with smooth boundaries along the nerve’s length, whereas the 2D U-Net produced jagged boundaries with evident artifacts (Figure 6c). We quantified this observation: along the longitudinal axis of the nerve, the 3D U-Net consistently demonstrated higher inter-slice consistency, maintaining BF score values above 0.8, which was a two-fold increase compared to the 2D U-Net (Figure 6d). On average, the 3D U-Net yielded significantly more consistent boundaries for both epineurium (0.77 vs. 0.60, *p* < 0.0001) and fascicles (0.81 vs. 0.77, *p* < 0.0001; Figure 6a). This boundary consistency is critical for tracking branches and fascicles along the nerve.

**Figure 6.**
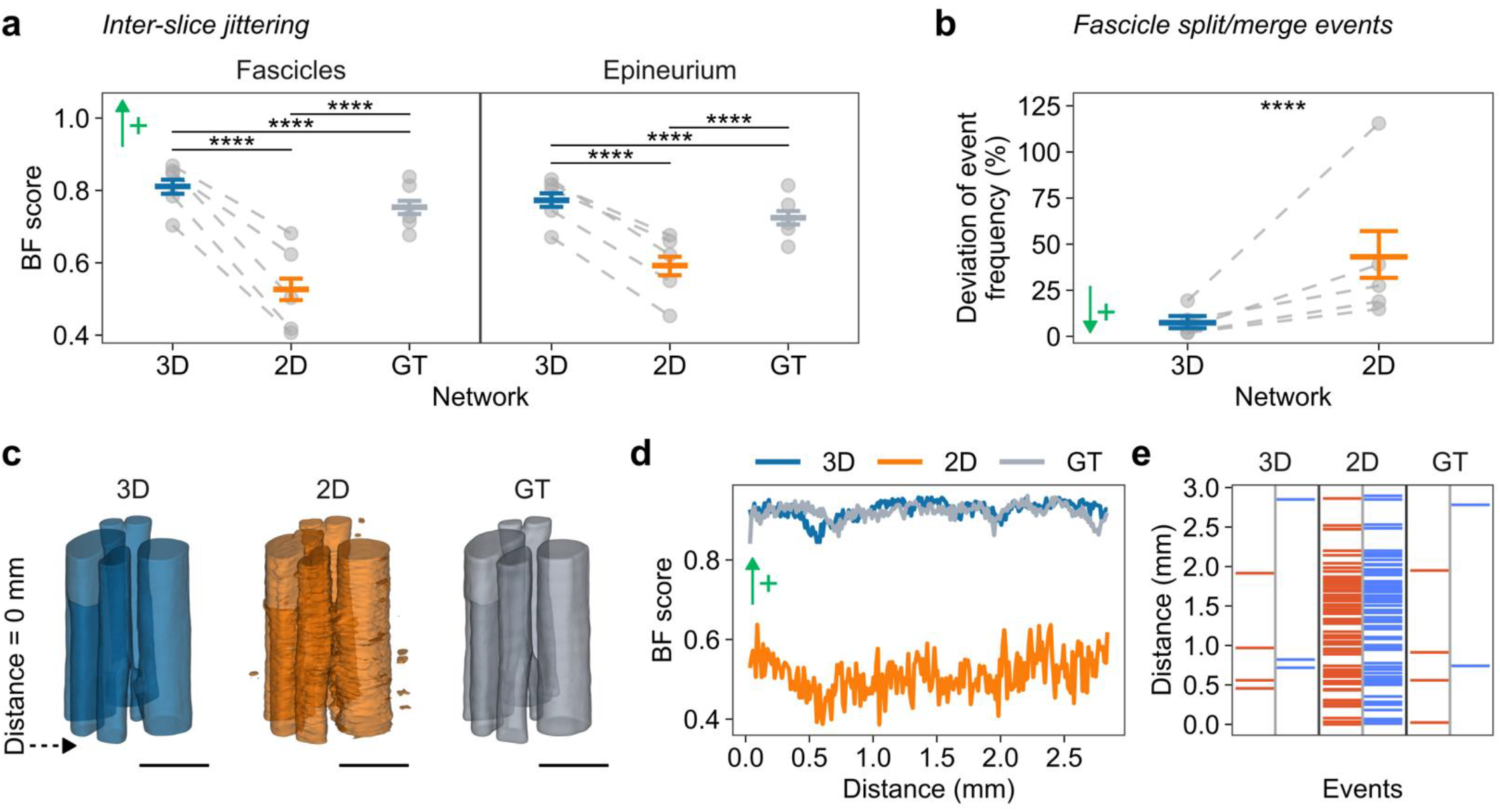
3D U-Net improves accuracy of longitudinal fascicular structure, including inter-slice jitter and split/merge events. **(a)** Boundary **F**_**1**_ (BF score) comparison for fascicles (left) and epineurium (right) across 3D/2D predictions and ground truth (GT). Higher scores indicate better boundary consistency between consecutive cross sections. **(b)** Fascicle split/merge event deviation from GT (%), measured as percentage difference between predicted and GT event frequency (number of splits and merges per millimeter). In **(a, b)**, plots show mean values with 95% confidence intervals; individual fold means are shown as gray dots connected by dashed lines. ↑+, higher is better; ↓+, lower is better. ****, *p* < 0.0001. **(c–e)** Example nerve sample showing: **(c)** 3D renderings of fascicle segmentations from 3D U-Net (blue), 2D U-Net (orange), and GT (gray); dashed arrow marks location of origins in (d, e). Scale bars are 1 mm. **(d)** BF score profiles along the example nerve sample in panel (c). **(e)** Distribution of fascicle merge (red) and split (blue) events for the example nerve sample in panel (c).

In the analysis of fascicle split/merge events, the 3D U-Net provided substantially better agreement with ground truth than the 2D U-Net (percentage deviation: 7.4% vs. 43.1%, *p <* 0.0001; Figure 6b). In contrast, the 2D U-Net generated a high number of false split/merge events, whereas the 3D U-Net correctly detected most true events (Figure 6e). When errors occurred, both networks tended to over-predict both split and merge events, resulting in a strong correlation between split and merge rate deviations (Figure S4 in Supplementary Note 7).

## 4 Discussion

We developed an enhanced deep learning approach for accurate, automated 3D segmentation of vagus nerve fascicular morphology from microCT images, designed to overcome limitations of existing techniques. Specifically, we used a 3D U-Net CNN to leverage volumetric context (Figure 1) and integrated a novel anatomy-aware loss function during training to enforce anatomical coherence (Figure 2). This methodology yielded substantial improvements compared to a 2D U-Net, as quantified by multiple evaluation metrics. Overall segmentation accuracy was significantly increased (Figure 3), and task-specific evaluations confirmed improved fascicle detection (Figure 4) and fewer anatomical errors (Figure 5). The proposed 3D U-Net also improved boundary continuity and more accurately identified fascicle splits/merges along the nerve (Figure 6). Collectively, these performance gains establish our method as a valuable tool for generating accurate, quantitative 3D anatomical maps of the vagal pathway from high-resolution imaging data. By addressing bottlenecks in segmentation quality and throughput, our approach provides data that are fundamental for understanding the nerve’s functional organization and for developing realistic computational models to guide VNS therapies.

### 4.1 3D U-Net Better Resolves Vagal Morphology than 2D U-Net

Segmenting the vagus nerve is challenging due to its complex morphology, with highly variable fascicle sizes and counts across individuals (Pelot et al., 2020) and along the nerve of a given individual (Upadhye et al., 2022). The longitudinal changes in vagal morphology expose a significant limitation of 2D segmentation methods (Buyukcelik et al., 2023; Jayaprakash et al., 2023), which process each slice independently and ignore inter-slice context. Here, we demonstrated the benefits of a 3D U-Net CNN for analyzing 3D morphology. This represents the first such application for peripheral nerve segmentation and extends the established success of volumetric deep learning for other intricate structures, such as vasculature (Merkow et al., 2016) and airways (Garcia-Uceda et al., 2021).

The 3D CNN’s ability to incorporate volumetric context resulted in superior segmentation performance. By integrating voxel information along the nerve, our 3D U-Net more accurately resolved the boundaries of fascicles and epineurium (Figure 3c–d). The 3D U-Net reduced the boundary artifacts (e.g., broken, spurious, or jagged edges) that are observed in the outputs of the 2D U-Net due to its slice-by-slice processing. Practically, the higher surface DSC achieved by the 3D U-Net will require less manual correction time following the automated segmentation (Figure 3c) (Vaassen et al., 2020).

Fascicle size and location are critical for computational modeling of peripheral nerve stimulation (Davis et al., 2023; Grinberg et al., 2008; Musselman et al., 2023). The 3D U-Net achieved higher fascicle-level accuracy within cross sections than the 2D U-Net (Figure 4b). With volumetric processing, the 3D segmentation mitigated incorrect merging of adjacent fascicles (i.e., under-segmentation), a critical and frequent failure mode in 2D methods that arises from imperfect contrast and small spacing between fascicles (Figure 4c). This accuracy gain in fascicle detection was size-dependent, with the most pronounced improvements observed for small- and medium-sized fascicles (0.16–0.62 mm in diameter) (Figure 4e), i.e., two-thirds of fascicles (Figure 4e Supplementary Note 5). Notably, both networks showed reduced performance in segmenting tiny fascicles (Figure 4e and Table S4 in Supplementary Note 5), likely due to the resolution limit of our microCT data; the insufficient definition of these smallest fascicles appeared to confound both manual annotation and network prediction, thus lowering the match rate.

### 4.2 Incorporating Anatomical Constraints Improves Segmentation Accuracy

Accurate segmentation of vagal morphology requires more than strong quantitative performance according to established generalized segmentation metrics: for meaningful downstream modeling and analysis, outputs must faithfully reproduce the anatomy of peripheral nerves. While U-Net architectures (Çiçek et al., 2016; Isensee et al., 2021; Ronneberger et al., 2015) provided a foundation for high-quality segmentation, we enhanced network training by incorporating structural priors, i.e., pre-existing knowledge about constraints on neuroanatomical structures. Specifically, we implemented an anatomy-aware loss function that supervised the network to learn the fundamental relationship wherein fascicles are enclosed by epineurium (Gupta et al., 2022; Pelot et al., 2020; Stewart, 2003). Our anatomy-aware loss function resulted in segmentations with improved anatomical integrity compared to baseline methods, preserving fascicle connectivity (Figure 5b) and reducing topological errors such as broken fascicles (Figure 5c). Our ablation study differentiated these effects, confirming that the anatomy-aware loss significantly minimized voxels violating structural rules, while the major gains in fascicle connectivity resulted from adopting the 3D architecture (Table S5 in Supplementary Note 6).

Prior studies also integrated anatomical knowledge in deep learning networks to improve segmentation beyond pixel classification (Gupta et al., 2022; Hu et al., 2019). This principle—implemented either through loss functions, network architectures, or postprocessing—has proven effective for structures with relatively predictable global topologies, such as tubular blood vessels (Shit et al., 2021), cardiac chambers (Oktay et al., 2018), and brain subcortical areas (Lorio et al., 2016). Our work extends this paradigm to peripheral nerve morphometry, suggesting that enforcing basic local spatial relationships of neural tissues can effectively regularize segmentation and improve anatomical fidelity for complex vagal structures.

### 4.3 Task-Specific Metrics Guide Meaningful Performance Assessment

In addition to our anatomy-aware loss function used for network training, we also implemented anatomy-aware metrics for post hoc evaluation of segmentation accuracy. Standard pixel-based segmentation metrics, such as DSC, often fail to reflect the practical utility of segmentation outputs (Müller et al., 2022). For instance, evaluating performance at object-level instead of pixel-level is a common approach for cell segmentation (Caicedo et al., 2019) and general computer vision (Kirillov et al., 2019), but its application specifically to peripheral nerve morphology has been limited. Only Buyukcelik *et al*. (2023) has reported fascicle-level accuracy. Our evaluation extended beyond geometric measures to assess factors essential for PNS modeling, neuroanatomical pathway mapping, and morphological measurements. Specifically, compared to a 2D U-Net, we demonstrated our 3D U-Net’s improved detection of fascicles of different sizes (Figure 4b and e), preserved fascicle connectivity (Figure 5b), reduced structural violations (Figure 5c), reduced discontinuities between transverse cross sections (Figure 6a), and a 6-fold decrease in errors detecting fascicle split/merge events (Figure 6b). This focus on task-specific performance is crucial because a segmentation with good DSC may still contain disconnected fascicular structures or abrupt cross-sectional changes, which hinder both longitudinal fascicle tracking and computational modeling.

Inter-slice jitter poses an important challenge to computational modeling of PNS, particularly for meshing the nerve geometry in a finite element model. We used the BF score to quantify inter-slice jitter, which is a metric that is sensitive to changes in contour connectivity (Csurka et al., 2013). However, the BF score decreases both when the network introduces discontinuities across slices and during true fascicle splitting and merging events. This ambiguity may complicate the specific assessment of network-induced errors (jitter) versus actual anatomical changes (splits/merges). Although we separately analyzed accuracy of splits and merges (Figure 6b, e), the BF score or an alternative metric could more definitively assess inter-slice jitter by applying it to only portions of fascicles without splits or merges.

### 4.4 Segmentation Accuracy Is Critical to Neuroanatomical Analyses and Computational Modeling

The accuracy of nerve morphology segmentation has important implications for anatomical analyses. Our automated 3D segmentation approach reduces boundary errors and enhances fascicle detection, thus providing a significantly more reliable source of morphometrics (e.g., fascicle count, size, spatial organization). This gain in accuracy is critical, as variations in morphology have important effects on neural responses to bioelectronic therapies (Davis et al., 2023; Grinberg et al., 2008). The ability to automatically extract precise morphology from high-resolution images enables analyses of anatomical variations within the vagus nerve across different locations, individuals, and species (Pelot et al., 2020). The 3D U-Net also stabilizes inter-slice boundaries, preserves fascicle connectivity, and accurately detects splitting and merging events, all of which are important for mapping the functional organization of peripheral nerves by proximally tracking fascicles that innervate specific organs and tissues. For example, histology and/or microCT have been used to map the functional organization of the pig vagus nerve (Jayaprakash et al., 2023; Settell et al., 2020; Thompson et al., 2025, 2023) and nerves of the upper limb in humans (Sunderland, 1945). However, fascicles in those nerves split and merge far less frequently than in the human vagus nerve. Precisely tracking fascicles to quantify the functional organization is therefore a foundational step in the development of selective VNS/PNS therapies.

The anatomical accuracy of nerve segmentations is crucial for computational modeling of PNS. Computational models enable the examination of mechanisms of action of neural stimulation therapies, as well as the design of electrode geometries, placement, and stimulation parameters (Ackermann et al., 2011; Aristovich et al., 2021; Romeni et al., 2020; Schiefer et al., 2012; Wongsarnpigoon and Grill, 2010). Computational models of PNS typically define the nerve morphology by extruding a single cross section, thus assuming constant cross-sectional morphology (Bucksot et al., 2021; Helmers et al., 2012; Musselman et al., 2023; Tebcherani et al., 2024). Conversely, a recent study developed a pipeline to model true 3D nerve morphology based on segmented microCT data (Marshall et al., 2025, in review), motivated by the observation that the morphology of human vagus nerves changes every ∼0.56 mm (Upadhye et al., 2022). Indeed, the use of microCT is becoming commonplace as a complement to histology to investigate peripheral neuroanatomy (Jayaprakash et al., 2023; Thompson et al., 2020; Upadhye et al., 2022). This convergence of advancements in computational modeling and imaging will benefit from segmentation algorithms that can robustly, accurately, and efficiently process imaging data into simulation inputs.

### 4.5 Application to Segmentation of Other Imaging Data

We expect our 3D U-Net segmentation approach to translate well to datasets with varied staining methods, nerve types, and species. For example, nerves are often stained with agents other than PTA for microCT imaging, such as osmium tetroxide (Upadhye et al., 2022) or Lugol’s iodine (Jayaprakash et al., 2023; Thompson et al., 2020); nerves other than vagus are also established targets for implanted neuromodulation devices, including the tibial nerve (Delianides et al., 2020; Rogers et al., 2021), sacral nerve (Li et al., 2016), hypoglossal nerve (Strollo et al., 2014), and median nerve (Tan et al., 2015); and animal models are often used for therapy development despite their different morphologies from humans (Pelot et al., 2020; Settell et al., 2020). In all cases, our anatomy-aware loss function and evaluation metrics can be applied directly, and our trained 3D U-Net may provide satisfactory results. However, in some cases—e.g., microCT images acquired with different stains or nerves with morphologies that differ substantially from human vagus nerves—the 3D U-Net may require either de novo training or transfer learning with dataset-specific ground truth annotations. Quantitative analyses of peripheral neuromodulation targets will benefit from our improved segmentation models and metrics.

Our 3D U-Net approach is more demanding than 2D U-Nets, requiring more time for generating ground truth annotations and computational resources for training and inference. Generating 3D ground truth annotations for training is more labor-intensive than 2D, requiring 30 minutes to 1.5 hours per 64-slice (0.73 mm-long) volume, while 2D annotation typically takes less than 5 minutes per cross section. Due to increased network parameters and image sizes, training and inference in 3D are generally slower than 2D on an equivalent device. For training and inference in 3D, memory constraints may require small batch sizes and patch-wise prediction, which can introduce the computational challenge of reconstructing the whole volume while ensuring the smooth blending of predicted patches. In the case of the human vagus nerve, the additional demands of the 3D U-Net are warranted given the frequent fascicle splits and merges (Upadhye et al., 2022), which are not adequately captured by a single or a stack of 2D images. The value of the 3D U-Net is demonstrated by its lower inter-slice jitter compared to both the 2D U-Net and ground truth (Figure 6a), in addition to overall higher segmentation quality and anatomical accuracy. For other nerves and objectives, it should be considered whether a 2D U-Net suffices.

## 5 Conclusion

We developed an enhanced deep learning approach, featuring a 3D U-Net CNN trained with an anatomy-aware loss function, for segmenting human vagus nerve fascicles and epineurium from microCT images. Compared to a 2D U-Net, our 3D U-Net provided a more reliable structural representation of the vagus nerve with improved segmentation accuracy, fascicle detection, and anatomical coherence. It also demonstrated enhanced performance in task-specific metrics critical to neuroanatomical analysis, including inter-slice consistency and accurate detection of fascicle branching events. These improvements enable more robust quantification of complex vagal morphology and provide high-fidelity 3D anatomical inputs essential for developing clinically relevant computational models of VNS. By automatically and accurately capturing the 3D vagus nerve morphology, this technique will provide the throughput required for large-scale characterization of intra- and inter-individual anatomical variability that influences VNS outcomes. Adaptable to other datasets, nerve targets, and species, this segmentation tool represents a critical advance within the broader strategy of leveraging high-resolution imaging and in silico modeling pipelines to support the rational design of precise, personalized neuromodulation therapies.

## 6 Acknowledgements

We thank the Department of Anatomy of Case Western Reserve University (Brandon Brunsman, Leina Lunasco, Andrew Crofton) for dissecting the human vagus nerve specimens. We thank the Department of Biostatistics & Bioinformatics at Duke University (Allison Yu, Sheng Luo) for support with statistical analyses. Funding was provided through the National Institute of Health (NIH) Stimulating Peripheral Activity to Relieve Conditions (SPARC) program (75N98022C00018), the US Department of Veterans Affairs (1IS1BX004384), Cleveland VA APT Center, and Case Western Reserve University. The opinions expressed in this paper are those of the authors and do not reflect the views of the NIH, the Department of Health and Human Services, or the United States government.

