## Supplemental for "Automated 3D Segmentation of Human Vagus Nerve Fascicles and Epineurium from Micro-Computed Tomography Images Using Anatomy-Aware Neural Networks"

**Supplementary Note 1 Human cadaver metadata.**

**Table S1.** Metadata for the human cadaveric subjects used in this study.

| Subject ID | Subject ID in Dataset | Sex | Age (years) | Self-reported race |
|---|---|---|---|---|
| 1 | SR007 | Male | 80 | White |
| 2 | SR008 | Female | 79 | White |
| 3 | SR009 | Male | 57 | Hispanic |
| 4 | SR010 | Female | 81 | White |
| 5 | SR011 | Male | 76 | N/A |

**Supplementary Note 2 Network training and 2D U-Net architecture.**

**Table S2.** Parameters and probabilities for data augmentation techniques applied during 3D and 2D network training.

| Augmentation | Parameters | Probability |
|---|---|---|
| Random rotation | Angle: $\pm 30$ ° for 3D, $\pm 180$ ° for 2D | 0.20 |
| Random scaling | Scale factor: 0.7–1.4 | 0.20 |
| Additive Gaussian noise | $\mu = 0$, noise variance: 0–0.1 | 0.15 |
| Gaussian blur | $\sigma$: 0.5–1.5 | 0.20 |
| Brightness adjustment | Factor: 0.7–1.3 | 0.15 |
| Contrast adjustment | Factor: 0.65–1.5 | 0.15 |
| Random downsampling | Downsampling factor: 1–2 | 0.25 |
| Gamma contrast transform | $\gamma$: 0.7–1.5 | 0.15 |
| Flipping | Along each axis | 0.50 |

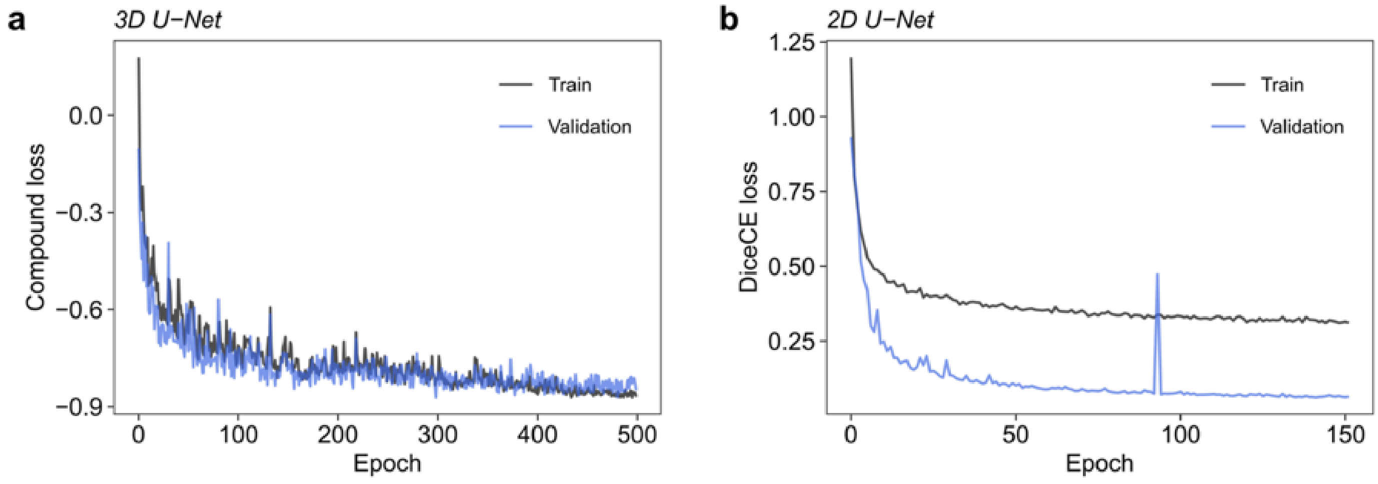


**Figure S1.** Example training and validation loss curves for the 3D and 2D U-Nets during the first cross-validation fold. Both networks were trained on data from subjects 2 to 5. **(a)** Compound loss for the 3D U-Net, trained and validated on randomly sampled patches (32 × 256 × 256) from full-resolution volumes (64 × 1536 × 3072). The compound loss represents a weighted sum of Dice, cross-entropy, and topological loss terms. **(b)** Dice and Cross Entropy (DiceCE) loss for the 2D U-Net, trained on randomly sampled patches (512 × 512) from full-resolution slices (1536 × 3072) and validated on whole slices.

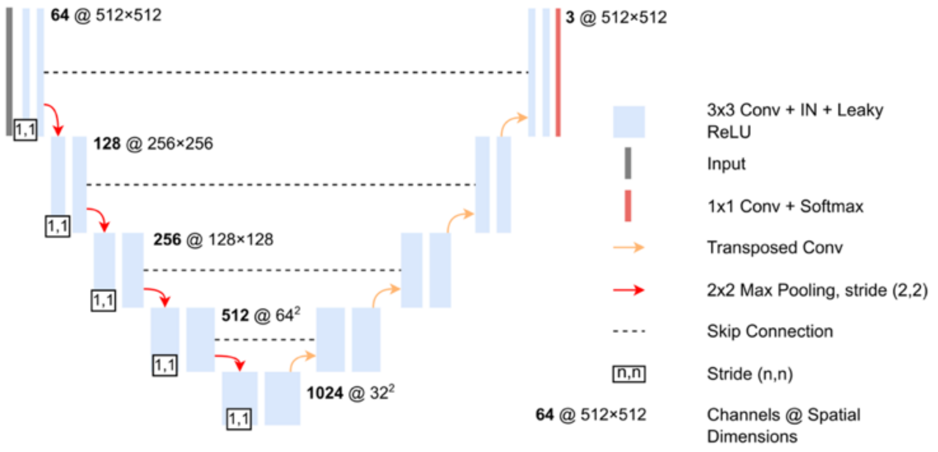


**Figure S2.** Architecture of the baseline 2D U-Net used for comparison to our 3D U-Net. Conv, convolution; IN, instance normalization; ReLU, rectified linear unit.

**Supplementary Note 3 Methodology for quantifying anatomical errors in network prediction.**

We measured the frequency of anatomical abnormalities in predicted vagus nerve segmentations by identifying structurally implausible voxels. Our method detects two categories of anatomical violations:

1. Exposed fascicle voxels that directly contact background.

   We identified exposed fascicle voxels ($V_{\text{exposed}}$) by:

   $$V_{\text{exposed}} = F \odot (B * K_{26})$$

   where $F$ is the binary fascicle mask, $B$ is the binary background mask, $\odot$ is the Hadamard product, and $K_{26}$ is the 26-connectivity kernel (a $3 \times 3 \times 3$ matrix of ones with zero at the center).

   $$K_{26}(i,j,k) = \begin{cases} 0, & \text{if } i = j = k = 1 \text{ (center position)} \\ 1, & \text{otherwise} \end{cases}$$

   The convolution $B * K_{26}$ identifies background-adjacent voxels, which when multiplied with $F$, yields fascicle voxels that directly contact the background.

2. Abrupt transitions between neural structures along the nerve.

   We identified abrupt transitions between neural structures along the nerve ($V_{\text{discontinuity}}$) by first identifying voxels where two different classes ($c, c'$) are present in adjacent slices:

   $$V_{c,c'} = (P_c * K_L > 0) \odot (P_{c'} * K_L > 0)$$

   where $P_c$ and $P_{c'}$ are the binary masks for classes $c$ and $c'$, respectively, and $K_L$ is the longitudinal kernel (a $1 \times 3 \times 3$ matrix of ones). This kernel structure is aligned with the nerve's longitudinal axis in all images.

   $$K_L(i,j,k) = \begin{cases} 1, & \text{if } j = k = 1 \text{ (central axis along the first dimension)} \\ 0, & \text{otherwise} \end{cases}$$

   To avoid counting normal structural boundaries, we excluded surface voxels from the class change maps:

   $$V_{c,c',\text{non-surface}} = V_{c,c'} \odot \neg(S_c \cup S_{c'})$$

   where $S_c$ and $S_{c'}$ are the sets of surface voxels for classes $c$ and $c'$, respectively. The overall class change map was the union of all class change maps for all class pairs:

   $$V_{\text{discontinuity}} = \bigcup_{\substack{c,c' \in C \\ c < c'}} V_{c,c',\text{non-surface}}$$

   where $C$ is the set of classes (i.e., fascicles, epineurium, and background).

Finally, the anatomical error rate was calculated as:

$$\text{Anatomical error rate} = \frac{|V_{\text{exposed}} \cup V_{\text{discontinuity}}|}{|V_{\text{foreground}}|} \times 100\%$$

where $V_{\text{foreground}}$ is the volume of the foreground mask (i.e., fascicles and epineurium).

**Supplementary Note 4 Additional segmentation metrics.**

**Table S3.** Comparison of voxel-based segmentation metrics for fascicles and epineurium using 3D and 2D U-Nets.

| | **Network** | | | | |
|---|---|---|---|---|---|
| **Metric** | **3D**, N = 100[a] | **2D**, N = 100[a] | **Difference**[b] | **95% CI**[b] | ***p*-value**[c] |
| Fascicles | | | | | |
| IoU | 0.8674 | 0.8280 | 0.03 | 0.01, 0.05 | <0.001 |
| Sensitivity | 0.9280 | 0.9123 | 0.01 | 0.00, 0.02 | 0.002 |
| Specificity | 0.9996 | 0.9993 | 0.00 | 0.00, 0.00 | <0.001 |
| Epineurium | | | | | |
| IoU | 0.8777 | 0.8544 | 0.02 | 0.01, 0.03 | <0.001 |
| Sensitivity | 0.9354 | 0.9204 | 0.01 | 0.00, 0.02 | 0.003 |
| Specificity | 0.9986 | 0.9986 | 0.00 | 0.00, 0.00 | 0.2 |

a. Mean
b. Wilcoxon rank sum test
c. Bonferroni correction for multiple testing

Abbreviations: CI = Confidence Interval; IoU = Intersection-over-union.

**Supplementary Note 5  Area-based fascicle detection analysis.**

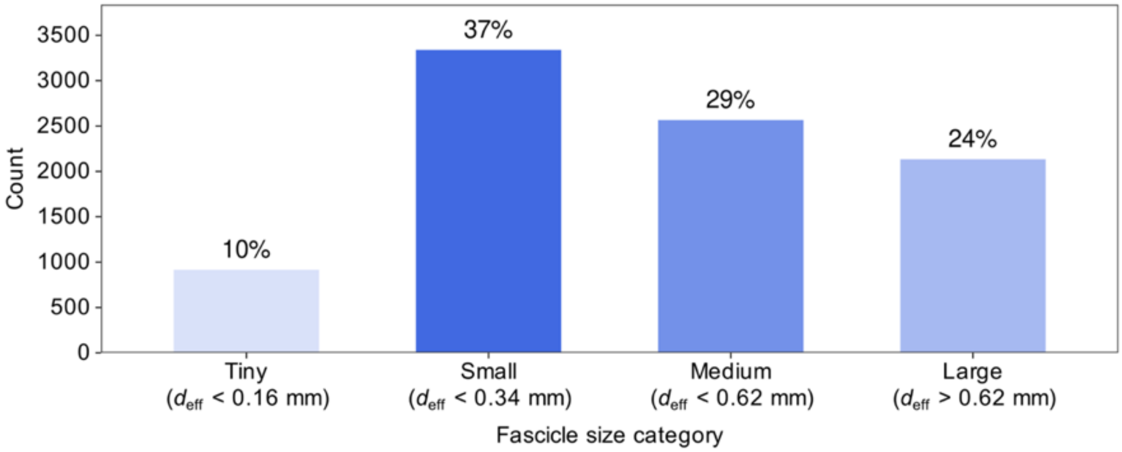


**Figure S3.** Distribution of vagus nerve fascicle areas grouped by size category in our microCT dataset. Fascicle size categories: tiny (area < 0.02 mm$^2$, $d_{eff}$ < 0.16 mm), small (area < 0.09 mm$^2$, $d_{eff}$ < 0.34 mm), medium (area < 0.3 mm$^2$, $d_{eff}$ < 0.62 mm), and large (area > 0.3 mm$^2$, $d_{eff}$ > 0.62 mm). $d_{eff}$, effective circular diameter.

**Table S4.** Comparison of instance segmentation metrics between 3D and 2D networks for matched human vagus nerve fascicle pairs. Matching was based on an intersection-over-union (IoU) threshold of 0.7. Metrics are presented across four fascicle area categories: tiny (<0.02 mm$^2$), small (0.02–0.09 mm$^2$), medium (0.09–0.3 mm$^2$), and large (>0.3 mm$^2$).

| | **Network** | | | | |
|---|---|---|---|---|---|
| **Metric** | **3D**[a] | **2D**[a] | **Difference**[b] | **95% CI**[b] | ***p*-value**[c] |
| | | | Tiny | | |
| IoU | 0.79 | 0.78 | 0.01 | -0.01, 0.03 | >0.99 |
| HD (μm) | 2.68 | 2.77 | -0.15 | -0.56, 0.25 | >0.99 |
| Area difference (%) | 16.66 | 19.16 | -2.84 | -6.50, 0.95 | 0.43 |
| | | | Small | | |
| IoU | 0.82 | 0.81 | 0.01 | 0.00, 0.01 | 0.001 |
| HD (μm) | 2.63 | 2.86 | -0.18 | -0.27, -0.08 | <0.001 |
| Area difference (%) | 12.09 | 12.20 | -0.19 | -0.71, 0.32 | >0.99 |
| | | | Medium | | |
| IoU | 0.86 | 0.85 | 0.01 | 0.01, 0.01 | <0.001 |
| HD (μm) | 2.91 | 3.19 | -0.22 | -0.31, -0.13 | <0.001 |
| Area difference (%) | 9.70 | 9.46 | 0.15 | -0.24, 0.54 | >0.99 |
| | | | Large | | |
| IoU | 0.90 | 0.88 | 0.01 | 0.01, 0.01 | <0.001 |
| HD (μm) | 6.82 | 6.26 | -0.21 | -0.31, -0.12 | <0.001 |
| Area difference (%) | 7.11 | 7.39 | -0.13 | -0.41, 0.14 | >0.99 |

a. Mean
b. Wilcoxon signed rank test
c. Bonferroni correction for multiple testing
d. Defined as $|A_{\text{pred}} - A_{\text{gt}}|/A_{\text{gt}} \times 100\%$ where $A_{\text{pred}}$ is the predicted area and $A_{\text{gt}}$ is the ground truth area.

Abbreviations: CI, confidence interval; IoU, intersection-over-union; HD, Hausdorff distance.

**Supplementary Note 6 Ablation study.**

**Table S5.** Ablation study results assessing the impact of the anatomical loss term on anatomical accuracy. The 3D Base network, identical in architecture to the full 3D network, was trained without the anatomical loss term. All other training parameters remained constant.

| | **Network** | | |
|---|---|---|---|
| **Metric** | **3D**, N = 100[a] | **3D Base**, N = 100[a] | ***p*-value**[b] |
| clDice | 0.90 | 0.89 | 0.2 |
| Error rate (%) | 0.65 | 0.77 | <0.001 |

a. Mean
b. Wilcoxon signed rank test with Bonferroni correction for multiple testing

**Supplementary Note 7 Correlation between split/merge rate deviations.**

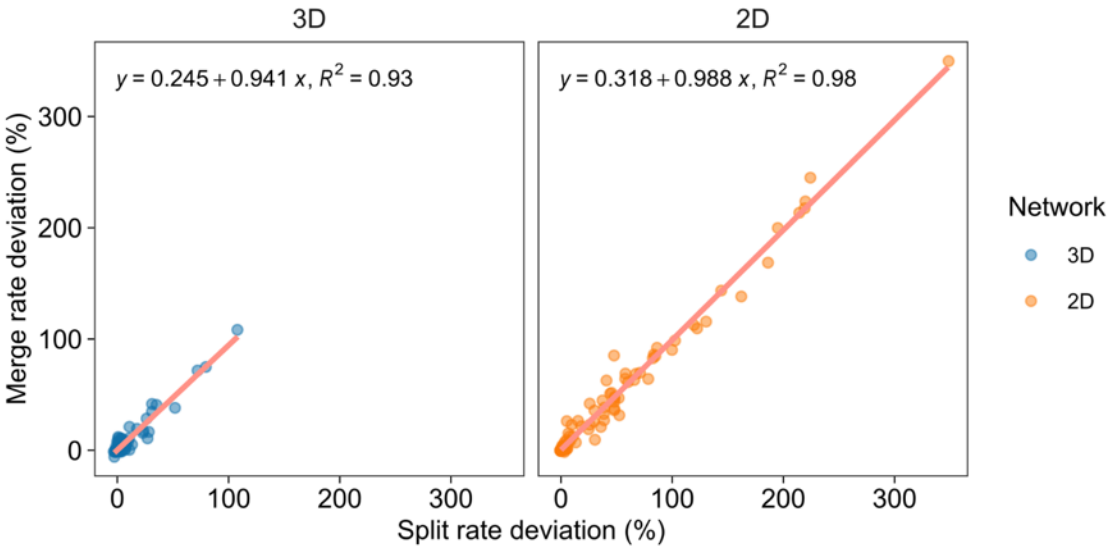


**Figure S4.** Correlation between split-rate and merge-rate deviations for the 3D and 2D networks. Deviation is defined as the percentage difference between the predicted and ground truth (GT) split/merge event frequency (number of split or merge events per millimeter).
